# Engineering Nitrogen Fixation Activity in an Oxygenic Phototroph

**DOI:** 10.1101/316547

**Authors:** Deng Liu, Michelle Liberton, Jingjie Yu, Himadri B. Pakrasi, Maitrayee Bhattacharyya-Pakrasi

**Author notes:** **Corresponding author**: Himadri Pakrasi, Department of Biology, Washington University, Campus Box 1137, One Brookings Drive, St. Louis, MO 63130.

## Abstract

Biological nitrogen fixation is catalyzed by nitrogenase, a complex metalloenzyme found only in prokaryotes. N_2_ fixation is energetically highly expensive, and an energy generating process such as photosynthesis can meet the energy demand of N_2_ fixation. However, synthesis and expression of nitrogenase is exquisitely sensitive to oxygen. Thus, engineering nitrogen fixation activity in photosynthetic organisms that produce oxygen is challenging. Cyanobacteria are oxygenic photosynthetic prokaryotes, and some of them also fix N_2_. Here, we demonstrate a feasible way to engineer nitrogenase activity in the non-diazotrophic cyanobacterium *Synechocystis* sp. PCC 6803 through the transfer of 35 nitrogen fixation (*nif*) genes from the diazotrophic cyanobacterium *Cyanothece* sp. ATCC 51142. In addition, we have identified the minimal *nif* cluster required for such activity in *Synechocystis* 6803. Moreover, nitrogenase activity was significantly improved by increasing the expression levels of *nif* genes. Importantly, the O_2_ tolerance of nitrogenase was enhanced by introduction of uptake hydrogenase genes, showing this to be a functional way to improve nitrogenase enzyme activity under micro-oxic conditions. To date, our efforts have resulted in engineered *Synechocystis* 6803 strains that remarkably have more than 30% N_2_-fixation activity compared to that in *Cyanothece* 51142, the highest such activity established in any non-diazotrophic oxygenic photosynthetic organism. This study establishes a baseline towards the ultimate goal of engineering nitrogen fixation ability in crop plants.

**IMPORTANCE:** Application of chemically synthesized nitrogen fertilizers has revolutionized agriculture. However, the energetic costs of such production processes as well as the wide spread application of fertilizers have raised serious environmental issues. A sustainable alternative is to endow crop plants the ability to fix atmospheric N_2_ *in situ*. One long-term approach is to transfer all *nif* genes from a prokaryote to plant cells, and express nitrogenase in an energy-producing organelle, chloroplast or mitochondrion. In this context, *Synechocystis* 6803, the non-diazotrophic cyanobacterium utilized in this study, provides a model chassis for rapid investigation of the necessary requirements to establish diazotrophy in an oxygenic phototroph.

---

Enabling crop plants the machinery to fix their own nitrogen via direct transfer of nitrogen fixation (*nif*) genes is envisioned to be key for the next agricultural revolution (1-3). However, engineering diazotrophic plants, attractive a proposition, will be an extreme challenge, due to the complexities in the biosynthesis of active nitrogenase, the enzyme that catalyzes nitrogen fixation, as well as the difficulty of coupling plant metabolism to supply energy and reducing power for the nitrogen fixation process (4). An additional impediment in the scenario is that photosynthesis produces O_2_, which is highly toxic to synthesis and activity of nitrogenase (5).

Diazotrophy occurs only in limited species of bacteria and archaea (6). Nitrogen fixation is mainly catalyzed by an iron and molybdenum-dependent nitrogenase enzyme complex, with two enzymatic components, an iron protein dinitrogenase reductase (NifH) and an iron-molybdenum protein dinitrogenase (NifDK) (7, 8). Three metal dependent cofactors, the F-cluster, P-cluster, and M-cluster, are necessary to form the holoenzyme for electron transfer to reduce atmospheric N_2_ to form ammonia, the biologically available form of N_2_ (9, 10). A significant number of additional *nif* genes are required for the biosynthesis of these metallocluster cofactors and for the maturation of nitrogenase to form a fully functional enzyme (11, 12).

Transferring nitrogen fixation to non-diazotrophs has been attempted for decades. To date, the heterotrophic bacterium *Escherichia coli* has been successfully engineered for nitrogen fixation activity through transfer of *nif* genes from various diazotrophic species (13-17). Engineering eukaryotic species for heterologous nitrogen fixation activity, including the yeast *Saccharomyces* cerevisiae and the green alga *Chlamydomonas reinhardtii*, have been unsuccessful. Limited success was reached only in expressing the NifH component as an active moiety in *Chlamydomonas reinhardtii* (18). While all the Nif components have been successfully expressed in yeast cells, the formation of a fully functional nitrogenase complex has not been achieved yet (19-22).

Expression of nitrogenase components into plants has also been attempted in a few studies. Individually expression of 16 Nif proteins targeted to the plant mitochondria has been reported recently, but none of the structural components showed enzymatic activity (23). Another recent study showed that an active NifH component can be formed in tobacco chloroplasts (24), indicating that expression of active nitrogenase in chloroplasts might be a viable way in the future to engineer crop plants to fix nitrogen(25). Since it is widely accepted that a cyanobacterial ancestor was the progenitor of chloroplasts (26), engineering a cyanobacterium to fix nitrogen may pave the way to achieve the final goal of engineering nitrogen fixing ability into crop plants. We have utilized the non-diazotrophic cyanobacterium *Synechocystis* sp. PCC 6803 (hereafter *Synechocystis* 6803) as a chassis to engineer nitrogen fixation activity into an oxygenic photosynthetic organism.

The unicellular diazotrophic cyanobacterium, *Cyanothece* sp. ATCC 51142 (hereafter *Cyanothece* 51142) uses temporal separation as its strategy to protect nitrogenase from O_2_ produced by photosynthesis (27, 28). Within *Cyanothece,* the two conflicting processes, photosynthesis and N_2_-fixation, occur sequentially during the diurnal periods, so that photosynthesis and O_2_ evolution is performed during the day whereas N_2_ is fixed at night (29). The energy requirements for nitrogenase are met in *Cyanothece* by the catabolism of glycogen. Glycogen is accumulated in the light as the storage form of fixed CO_2_, and later degraded in the dark to provide energy for nitrogenase. The provision of energy coupled with high rates of respiration ensures a low-oxygen intracellular environment and sufficient supplies of energy for N_2_ fixation (30). The *Cyanothece* 51142 genome contains the most complete contiguous set of nitrogen fixation and related genes to form a *nif* cluster, which contains 35 genes (cce_0545 to cce_0579), encoding structural proteins, metal cofactor synthesis proteins, ferredoxins, and genes for proteins with unknown but necessary functions (Fig. 1). All 35 genes exhibit similar oscillating diurnal pattern of transcription during light/dark cycles, showing a high level of transcription in the dark and notably reduced levels in light (27). Such a synchronized transcriptional pattern also confirms that all of these genes are related to nitrogen fixation.

**FIGURE 1.**
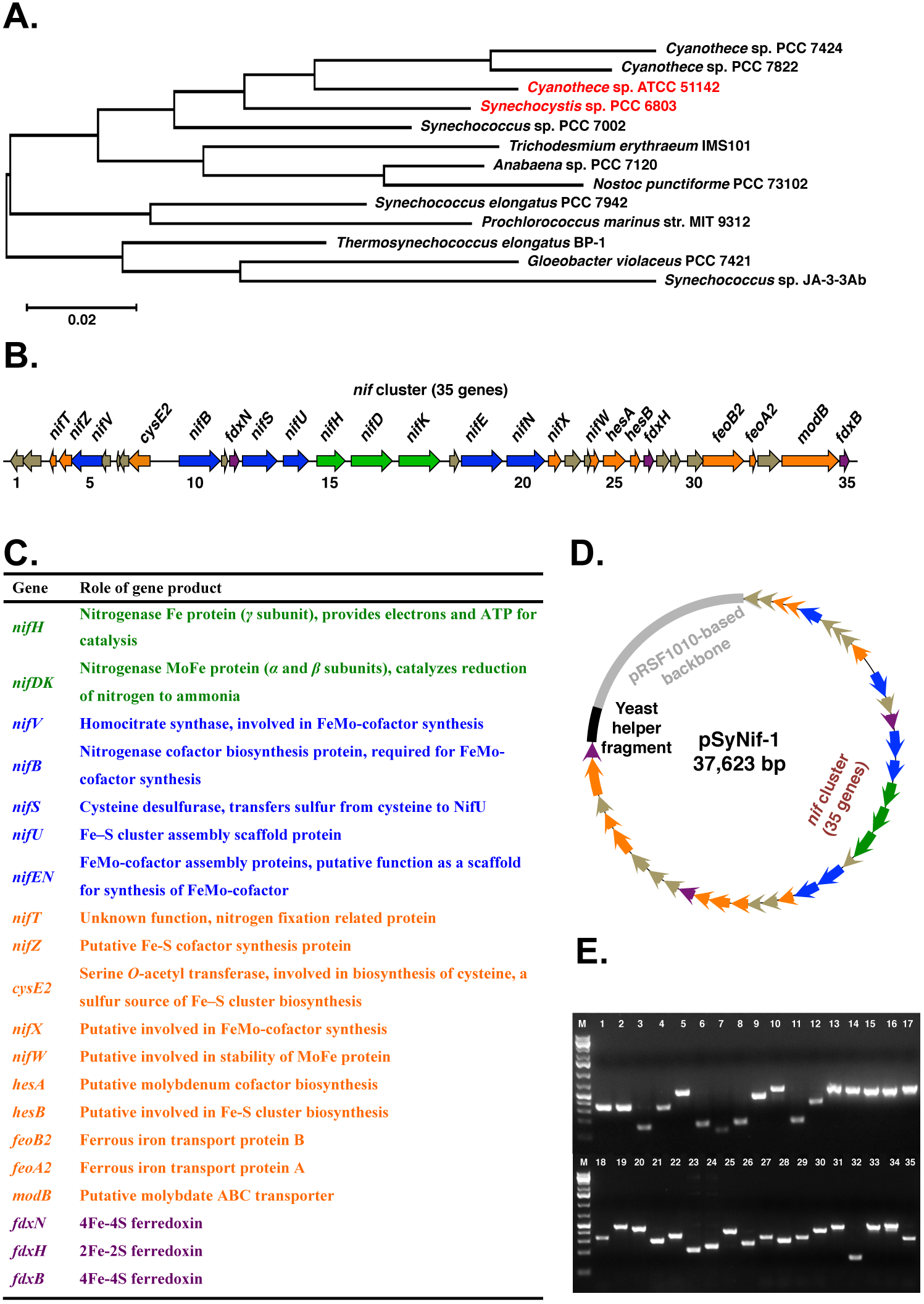
Introduction of *nif* genes into *Synechocystis* 6803. (A) Maximum likelihood 16S rDNA phylogeny of cyanobacteria. (B) Scheme showing the genetic organization of the *nif* cluster and (C) the role of each gene product in *Cyanothece* 51142. Shown are the genes for the three structural proteins, *nifHDK* (green), necessary cofactors (blue), accessory proteins (orange), ferredoxins (purple), and hypothetical proteins (brown). Gene names and annotation are from Genbank (https://www.ncbi.nlm.nih.gov/genbank/) and Cyanobase (http://genome.microbedb.jp/cyanobase). (D) A schematic map of the plasmid pSyNif-1 containing the entire *nif* cluster. The backbone (gray) is from the broad host plasmid pRSF1010, which can replicate in *Synechocystis* 6803. The yeast helper fragment (black) contains *CEN6* and *ARS* as an *ori*, and *ura3* as a selection marker. (E) Transcription of all 35 genes in engineered *Synechocystis* 6803. Each lane represents a gene in the *nif* cluster, as numbered in panel B. Total RNA was extracted from cells cultured in BG11_0_ medium under 12h light/dark conditions, and cDNA was used as the template for PCR.

In the current study, we have successfully transferred and expressed this large *nif* gene cluster in non-diazotrophic *Synechocystis* 6803 with a resultant N_2_-fixation activity. Subsequent engineering of the cluster as well its expression levels have led to nitrogenase activities as high as 30% of that in *Cyanothece* 51142.

## RESULTS AND DISCUSSION

### Introduction of *nif* genes into *Synechocystis* 6803

*Synechocystis* 6803 has a close phylogenetic relationship with *Cyanothece* 51142 (Fig. 1A) (31). The large *nif* cluster from *Cyanothece* 51142 (28.34 kilobase region of DNA) was transferred into wild type *Synechocystis* 6803 on a single extra-chromosomal plasmid. This large plasmid pSyNif-1 containing the entire *nif* cluster with 35 genes (Fig. 1D) was constructed using the DNA assembler method (32). The chassis of this vector was based on pRSF1010 (33), and this self-replicating plasmid pSyNif-1 was transferred into *Synechocystis* 6803 by conjugation, generating the engineered strain TSyNif-1. Remarkably, over the past four years since its introduction into the heterologous host, pSyNif-1 has been stably maintained in its entirety in *Synechocystis* (Fig. S1). Furthermore, all introduced *Cyanothece* genes were transcribed (Fig. 1E) as detected by RT-PCR, indicating that native promoters in the *nif* cluster from *Cyanothece* 51142 can drive transcription of genes in *Synechocystis* 6803. An acetylene reduction assay method for nitrogen fixation detected nitrogenase activity, under 12h/12h light/dark conditions for strain TSyNif-1 (Fig. 2). Nitrogen fixation reached an activity of 2% relative to that in *Cyanothece* 51142 grown under similar conditions. This is the first time that a non-diazotrophic phototroph has been engineered for biosynthesis of a fully functional nitrogenase enzyme and exhibits detectible and stable nitrogen fixation activity.

**FIGURE 2.**
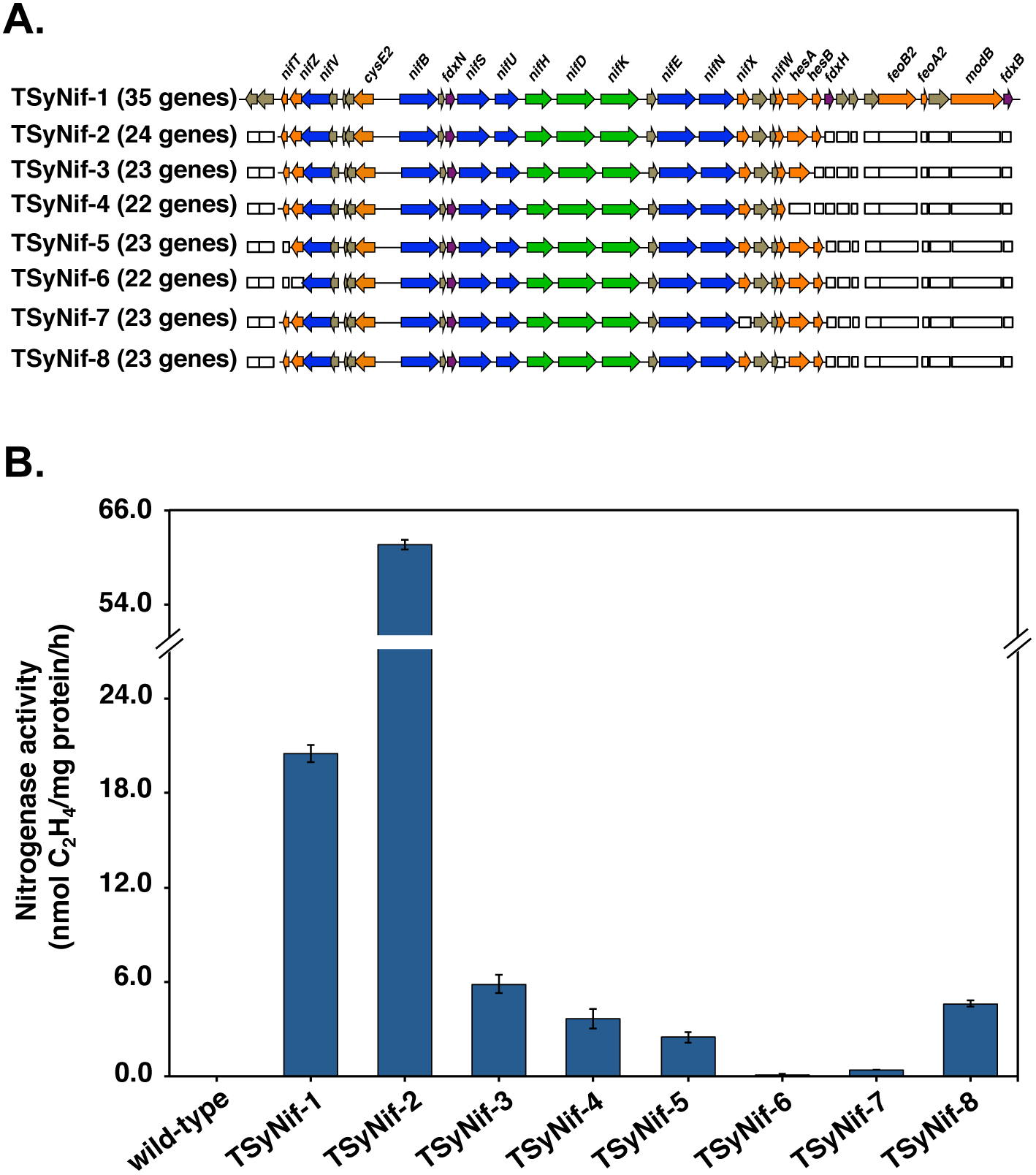
The minimal *nif* cluster required for nitrogen fixation activity in *Synechocystis* 6803. (A) Scheme showing the top-down method to determine the minimal *nif* cluster. The hollow rectangles represent the genes deleted from the cluster, and the colored rectangles represent the remaining genes. (B) Nitrogen fixation activity in engineered strains. Samples were collected from cultures under 12h light/dark conditions in BG11_0_ medium. Nitrogen fixation activity was assayed by acetylene reduction, and error bars represent the standard deviation observed from at least three independent experiments.

### The minimal required gene cluster for nitrogen fixation activity

Gene expression parameters for *Synechocystis* 6803 are not as well understood as for *E. coli*. Thus, the refactoring of *nif* genes as performed in *E. coli* and *Klebsiella* (16, 34) to determine the minimal requirement of genes for nitrogen fixation in *Synechocystis* 6803 is impractical at this stage. Therefore, we approached the identification of a minimal *nif* cluster for nitrogen fixation using a “top-down” method, which determines the influence of a gene on nitrogenase activity by selectively removing individual genes from the *nif* cluster (Fig. 2). Extrapolating the genetic requirements for nitrogen fixation activity observed in studies in which *nif* genes were introduced in *E. coli* (14, 15), we determined that genes for all homologous proteins introduced into *E. coli* are present in the *Cyanothece* 51142 *nif* cluster between gene *nifT* and *hesB* (Fig. 1). Hence, our second plasmid, pSyNif-2, contains 24 genes in the *nif* cluster between *nifT* and *hesB* (Fig. 2). Eleven genes were removed that presumably encode three metal transporter proteins, two ferredoxins, and six proteins of unknown function, none of which have been analyzed previously, although they are associated with nitrogen fixation. Intriguingly, this second engineered strain TSyNif-2 with a reduced cluster of 24 genes has a 3-fold increase in nitrogen fixation activity when compared to strain TSyNif-1 (Fig. 2). Although plasmids pSyNif-1 and pSyNif-2 have the same plasmid backbone (Fig. S2), the transcriptional levels of the structural genes *nifH, nifD* and *nifK* are higher in the TSyNif-2 strain (Fig. 3). This improvement in nitrogenase activity could be the result of removal of one or more regulatory gene(s), which may encode protein(s) that repress expression of genes in the *nif* cluster.

**FIGURE 3.**
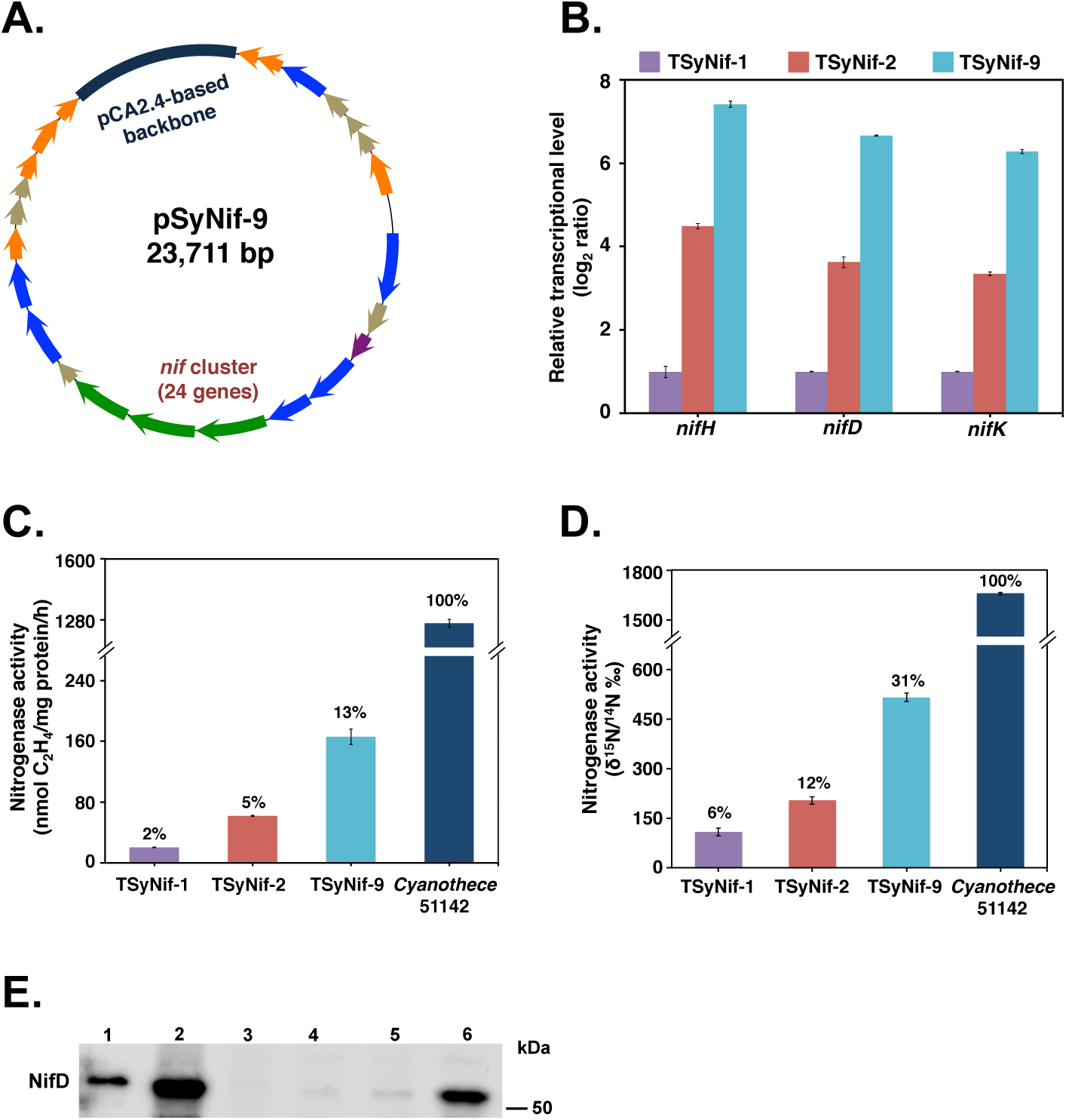
Enhancement of transcription levels of *nif* genes leads to higher nitrogen fixation activity. (A) A schematic map of the plasmid pSyNif-9 containing the *nif* cluster with 24 genes from *nifT* to *hesB*. The backbone (dark blue) is from the endogenous plasmid pCA2.4 of *Synechocystis* 6803. (B) Comparison of transcription levels of the *nif* structural genes in engineered strains through quantitative PCR (q-PCR). (C and D) Comparison of nitrogen fixation activities in engineered strains, as measured by (C) C_2_H_2_ reduction assay as well as (D) ^15^N assimilation assay. (E) Western blot showing the presence of NifD protein in engineered *Synechocystis* 6803 strains. Lanes 1-6 represent 0.5 μg purified NifD-His protein from *E. coli*, 15 μg whole cell extracts of *Cyanothece* 51142, *Synechocystis* 6803 wild-type, TSyNif-1, TSyNif-2, and TSyNif-9, respectively. Cyanobacterial samples were collected from cultures under 12h light/dark conditions in BG11_0_ medium. Error bars represent the standard deviation of at least three independent experiments.

However, further removal of genes from both directions, resulted in a decrease of nitrogen fixation activity by more than ten-fold for strains TSyNif-3 to TSyNif-6, in which the genes *hesB*, *hesAB*, *nifT*, and *nifTZ*, were removed, respectively (Fig. 2). Thus, this “top-down” approach determined that an essential minimal cluster from *nifT* to *hesB* is required for nitrogen fixation activity in *Synechocystis* 6803. Additionally, we investigated the removal of two more genes in the cluster, *nifX* and *nifW*, generating the two strains TSyNif-7 and TSyNif-8. Deletion of these two genes did not affect expression of surrounding genes (Fig. S3). However, nitrogen fixation activity dropped 100-fold and 10-fold, respectively, in these engineered strains (Fig. 2). We conclude that both *nifX* and *nifW* are important genes for nitrogen fixation. Notably, *nifX* exhibited a positive influence on N_2_ fixation in cyanobacteria, while it functions as a negative regulator for N_2_ fixation in the heterotrophic diazotroph *Klebsiella oxytoca* (35).

### Improvement of nitrogen fixation activity

To increase the RNA expression levels of the nitrogenase related genes, we took advantage of three small endogenous plasmids in *Synechocystis* 6803, pCA2.4, pCB2.4, and pCC5.2. The expression of heterologous genes expressed from these endogenous plasmids maintains higher transcriptional levels than from an pRSF1010 based plasmid, because of the higher copy numbers of these three plasmids within *Synechocystis* (36, 37). First, we replaced the RSF1010 backbone of plasmid pSyNif-2 by the entire DNA segment of each of these endogenous episomes and then transferred the plasmids to *Synechocystis* 6803, generating three strains TSyNif-9, TSyNif-10, and TSyNif-11, with the chassis of pCA2.4, pCB2.4, and pCC5.2, respectively (Fig. 3A and Fig. S4). As expected, the expression levels of genes *nifH*, *nifD* and *nif*K in strain TSyNif-9 showed several fold higher transcription levels compared to in TSyNif-2 (Fig. 3B). In addition, nitrogen fixation activity was increased by another 2 to 3-fold, reaching 13% of the acetylene reduction activity in TSyNif-9 relative to that observed in *Cyanothece* 51142 (Fig. 3C). Next, nitrogenase activity was directly assayed in the engineered strains using a 15N assimilation assay method. Remarkably, the highest activity obtained was from strain TSyNif-9, reaching 31% of 15N assimilation relative to *Cyanothece* 51142 (Fig. 3D). The activity data presented here is comparable to published data from studies on nitrogen fixation activity in engineered in *E. coli* (Table 1). Additionally, the dinitrogenase protein NifD in whole cell extracts was detected via Western blotting by using antisera from *Rhodospirillum rubrum* (Fig. 3E). Although the MoFe-protein level in *Cyanothece* 51142 is significantly higher than in the engineered *Synechocystis* 6803 strains, the protein level in strain TSyNif-9 reached 10% of total cellular proteins (Fig. S4). It was also evident that the nitrogenase activities in the engineered strains were proportional to the level of nitrogenase structural proteins, which implied that optimizing the expression of nitrogenase proteins is critical for the activity. Most importantly from an evolutionary standpoint, these results highlight the potential for engineering plant chloroplasts to fix nitrogen at a high level of activity, since oxygenic cyanobacteria are the progenitors of chloroplasts.

**Table 1.** Nitrogen fixation activity in diazotrophs and engineered strains.

| Strain | Nitrogenase activity<br>(nmol C<sub>2</sub>H<sub>4</sub>/mg protein/h) | % activity<br>(based on C<sub>2</sub>H<sub>2</sub> reduction assay) | % activity<br>(based on <sup>15</sup>N assimilation assay) | Reference |
| --- | --- | --- | --- | --- |
| **Diazotrophs** |  |  |  |  |
| *Cyanothece* 51142 | 1262 | 100 | 100 | This study |
| *Azotobacter vinelandii* | 3300 | 100 | 100 | (52) |
| *Paenibacillus* sp. WLY78 | 3050 | 100 | 100 | (14) |
| *Pseudomonas stutzeri* A1501 | 1050 | 100 | ND | (53) |
| *Klebsiella oxytoca* M5a1 | 3708 | 100 | ND | (54) |
| **Engineered strains** |  |  |  |  |
| TSyNif-1 | 20 | 2 | 6 | This study |
| TSyNif-9 | 166 | 13 | 31 | This study |
| Engineered *E. coli*<sup>a</sup> | 180 | 5 | 35 | (15) |
| Engineered *E. coli*<sup>b</sup> | 300 | 10 | 30 | (14) |
| Engineered *E. coli*<sup>c</sup> | 105 | 10 | ND | (53) |
| Engineered *E. coli*<sup>d</sup> | 740 | 20 | ND | (16) |

<sup>a</sup>Nitrogen fixation genes from *A. vinelandii* and *K. oxytoca*.
<sup>b</sup>Nitrogen fixation genes from *Paenibacillus*.
<sup>c</sup>Nitrogen fixation genes from *Pseudomonas*.
<sup>d</sup>Nitrogen fixation genes from *K. oxytoca*.
ND-Not determined

### Nitrogen fixation activity in *Synechocystis* 6803

Despite an additional metabolic load of expressing large cohorts of 35 or 24 genes related to nitrogen fixation being introduced in *Synechocystis* 6803, remarkably, the expression and activities of these heterologous proteins did not affect the growth of the engineered strains under diurnal light/dark conditions (Fig. S1). We used strain TSyNif-2 to assess the influence of oxygen and exogenous nitrate on nitrogenase activity under four conditions, BG11, BG11_0_ (BG11 without nitrate), BG11 with 10 μM DCMU (no O_2_ evolution) and BG11_0_ with 10 μM DCMU (Fig. S5). Interestingly, transcript levels of *nifH*, *nifD* and *nifK* genes in the TSyNif-2 strain were downregulated by nitrate, which is similar to that in *Cyanothece* 51142. Specifically, the depletion of nitrate improved the nitrogenase activity over 30-fold in BG11_0_ with DCMU (Fig. S5). Nitrogen fixation activity was obtained only in anaerobic environment when DCMU was added to the testing culture, although the headspace of all cultures was flushed with pure argon. These data indicate that oxygen generated by photosynthesis directly blocks nitrogenase activity in TSyNif-2, highlighting that one of the biggest challenges for engineering nitrogen fixation in oxygenic phototrophs is the sensitivity of nitrogenase to oxygen.

### Improvement of O_2_ tolerance by introduction of uptake hydrogenase

In order to test the oxygen sensitivity of nitrogen fixation activity in TSyNif-2, a measured amount of oxygen was added to the headspace to cultures grown in BG11_0_ media, to generate micro-aerobic conditions of 0.5% and 1.0% of O_2_ in the sealed testing bottles. The activity dropped more than 10-fold and 60-fold (Fig. 4A), respectively, demonstrating that as expected, nitrogen fixation activity in engineered *Synechocystis* 6803 is highly sensitive to O_2_. To enhance O_2_ tolerance under these same conditions, genes coding for the uptake hydrogenase enzyme from *Cyanothece* 51142 were introduced into the chromosome of the TSyNif-2 strain. The uptake hydrogenase is conserved in diazotrophic cyanobacteria (38), and has been shown to be necessary for nitrogen fixation under aerobic conditions in *Cyanothece* (39). The structural genes *hupS* and *hupL*, present together in a single operon in *Cyanothece* 51142, were transformed into TSyNif-2, generating strain TSyNif-12 (Fig. 4B). In addition to the structural genes *hupSL*, a protease encoded by *hupW* is also present in *Cyanothece* 51142. HupW is required for the maturation of HupL protein through the processing of its C-terminus (40). So, the *hupSLW* genes organized in two operons were transformed into TSynif-2 to generate the TSyNif-13 strain (Fig. 4B). The expression of *hup* genes in TSyNif-12 and TSyNif-13 was assessed by RT-PCR (Fig. S6). The introduction of the uptake hydrogenase did not affect nitrogen fixation activity under anaerobic conditions (Fig. 4C). Interestingly, under micro-aerobic conditions, nitrogen fixation activity markedly improved with the expression of uptake hydrogenase, especially for strain TSyNif-13, a 2-fold and a 6-fold increase in TSyNif-2 for O_2_ levels of 0.5% and 1.0%, respectively. The above results suggest that expression of uptake hydrogenase proves to be highly effective in enhancing O_2_ tolerance of nitrogen fixation activity in the engineered *Synechocystis* strain.

**FIGURE 4.**
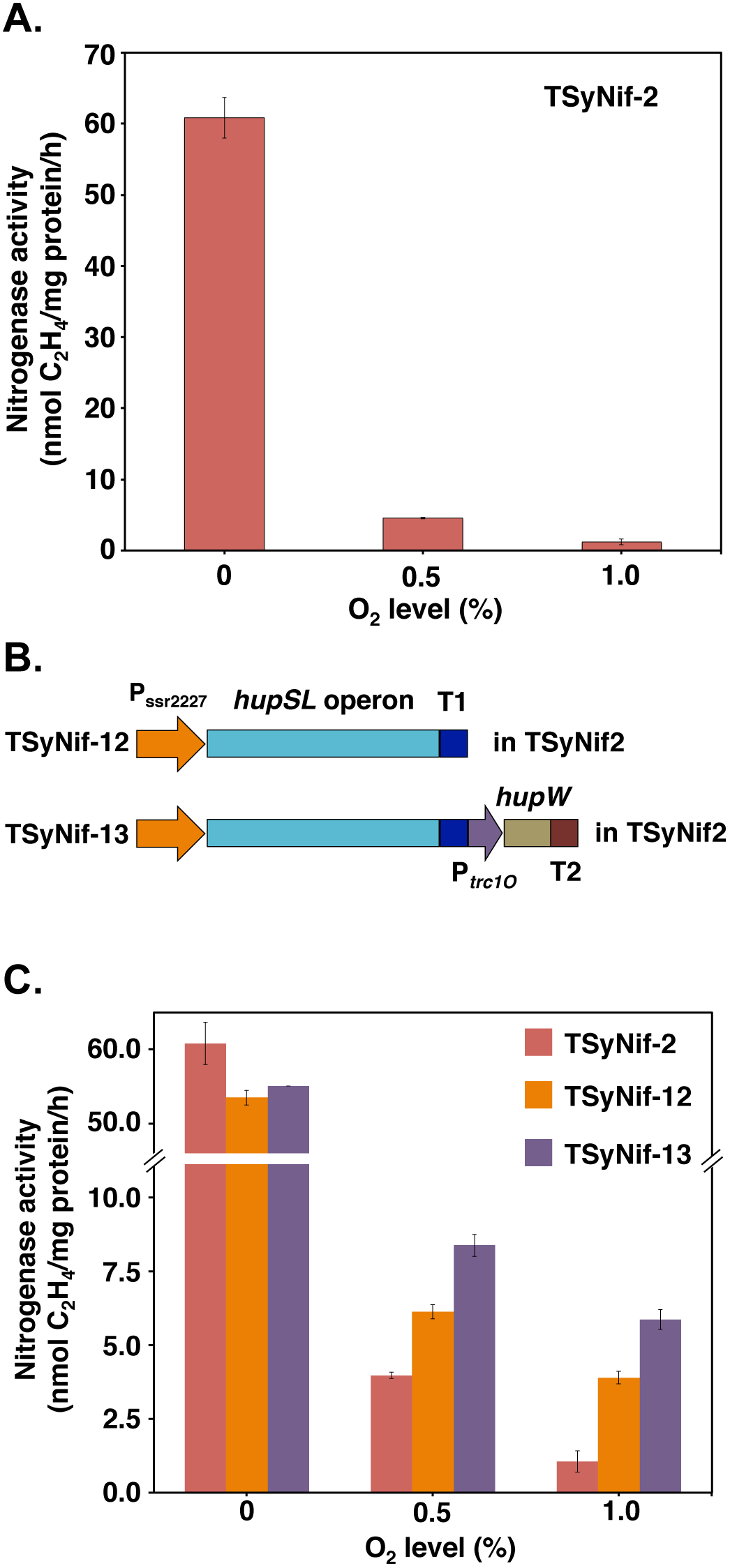
Expression of uptake hydrogenase improves O_2_ tolerance of nitrogenase. (A) Effect of O_2_ on nitrogen fixation activity of the TSyNif-2 strain. (B) Schematic showing the insertion of uptake hydrogenase genes *hupSL* and *hupW* from *Cyanothece* 51142 into the chromosome of the TSyNif-2 strain. (C) Comparison of nitrogen fixation activity under different micro-oxic conditions. Samples were collected from cultures under 12h light/dark conditions in BG11_0_ medium. Nitrogen fixation activity was assayed by acetylene reduction, and error bars represent the standard deviation of at least three independent experiments.

In this study, introduction of the *nif* gene cluster from *Cyanothece* 51142 has enabled nitrogen fixation activity in *Synechocystis* 6803. The minimal cluster for 24 genes (Fig. 2) for nitrogenase activity will provide a useful framework for future studies to further enhance such activity by refactoring genes as done in *Klebsiella oxytoca* (34). Although uptake hydrogenase is a complex enzyme, introduction of its structural genes and a protease works as a protecting mechanism from the toxicity of O_2_. The O_2_ toxicity to nitrogenase is likely the most difficult aspect to overcome to achieve nitrogen fixation activity under aerobic conditions.

A fully functional nitrogenase holoenzyme, requires 8 electrons and 16 ATPs to reduce one molecule of N_2_ to ammonia. Thus, metabolism within cells needs to be adjusted to supply enough reducing power and energy for nitrogen fixation. Biosynthesis of fully functional nitrogenase is a complex process. This complexity increases the difficulty to find the minimal genes and best ratios between each protein expressed from the *nif* cluster in *Synechocystis* 6803. The same gene designations in different species occasionally have alternative functions (Table S1). An example is the gene *nifX*, which functions as a negative regulator in *Klebsiella oxytoca (35)*. Gene *nifX* is of importance in *Cyanothece*, since deletion of *nifX* affects nitrogenase activity.

Although multiple challenges and many barriers exist to enable plants to efficiently fix atmospheric nitrogen, we have engineered an oxygenic photosynthetic cell to fix N_2_ by reconfiguring the genetic processes for nitrogen fixation from *Cyanothece* 51142 to function in *Synechocystis* 6803. Our studies to date have established the highest rate of engineered nitrogen fixation activity in any non-diazotrophic oxygenic organism.

## MATERIALS AND METHODS

### Microorganisms, culture conditions and media

All cyanobacterial strains, including *Cyanothece* 51142, *Synechocystis* 6803, and the engineered strains (listed in Table S2), were cultured in 100-mL flasks of fresh BG11 medium (41) with appropriate antibiotics (20 μg/ml kanamycin, 15 μg/ml chloramphenicol, or 20 μg/ml spectinomycin). As a pre-culture, cells were grown at 30°C, with 150 rpm shaking, and under 50 μmol photons · m^−2^ · s^−1^ constant light. For the nitrogen fixation activity assay, unless otherwise stated, pre-cultured cells were collected and washed with fresh BG11 without nitrate (BG11_0_) medium and resuspended in 500 ml fresh BG11_0_ medium. Cells were then grown at 30°C with air bubbling under 12h light/dark conditions with 50 μmol photons · m^−2^· s^−1^ of light. Yeast and *E. coli* strains (listed in Table S2) used for construction of recombinant plasmids were grown with 200 rpm shaking in YPAD (42) and LB medium at 30°C and 37°C, respectively.

### Construction of recombinant plasmids and engineered strains

Plasmids and strains used in this study are listed in Table S2, and all primers are listed in Table S3. Two methods were used to construct the plasmids, DNA assembler (32) and Gibson assembly (43). To build the large pSyNif-1 and pSyNif-2 plasmids containing the *nif* genes, genomic DNA from *Cyanothece* 51142 was used as the template for PCR, and all DNA fragments were combined using the DNA assembler method to construct the plasmids in the yeast *S. cerevisiae*. For the other recombinant plasmids listed in Table S2, Gibson assembly method was used to construct them with DNA fragments amplified by PCR. Genomic DNA from *Cyanothece* 51142 and the large plasmid pSyNif-2 were used as templates for PCR to construct the plasmids for backbone replacements, the plasmids containing the uptake hydrogenase genes, and the plasmids used to remove specific *nif* genes, respectively.

Plasmids pSyNif-1 and pSyNif-2 were introduced into *Synechocystis* 6803 wild-type strain through the method of tri-parental conjugation (44) to form strains TSyNif-1 and TSyNif-2, respectively. The other recombinant plasmids were transformed into *Synechocystis* 6803 by natural transformation (45), and double homologous recombination integrated the fragments into the chromosome (for the uptake hydrogenase genes) or the plasmid (for the plasmid backbone replacement, and specific removal of *nif* genes).

### Reverse transcription-PCR (RT-PCR) and Quantitative-PCR (q-PCR)

RT-PCR analysis was performed using RNA samples isolated from culture grown in BG11_0_ medium at time point D1 (1 hour into the dark period) under light/dark conditions. After extraction and quantification of RNA (46), 100 ng of DNase-treated RNA samples were used for reverse transcription, using the Superscript II Reverse Transcriptase and random primers (Invitrogen) according to the manufacturer’s instructions. cDNA generated after reverse transcription was used as the template for PCR to evaluate the transcription levels of various individual genes.

The q-PCR was performed on RNA samples extracted from culture grown in BG11_0_ medium under light/dark conditions as previously described (27). Briefly, QRT-PCR SYBR green dUTP mix (Abgene) was used for the assay on an ABI 7500 system (Applied Biosystems). Each reaction was performed in three replicates, and the average C_T_ was used to calculate the relative transcriptional levels for the amount of RNA. All primers used for RT-PCR and q-PCR are listed in Table S3.

### Measurement of nitrogen fixation activity

Nitrogen fixation activity was measured by an acetylene reduction assay (47), modified from a previously published method (48). Unless otherwise stated, the activity assay was performed as follows: 25 ml of cyanobacterial culture were grown in BG11_0_ medium with air bubbling under light/dark conditions as mentioned above, and transferred to a 125 ml air-tight glass vial. 10 μM DCMU was added to the culture, vials were flushed with pure argon, and cultures were incubated in 12h light/dark conditions. Cells in the sealed vials were cultured overnight and at the time point D1, 5 ml acetylene was added into the sealed vials, followed by 3 hours of incubation in light at 30°C. 200 μl of gas was sampled from the headspace and injected into an Agilent 6890N Gas Chromatograph equipped with a Poropak N column and a flame ionization detector, using argon as the carrier gas. The temperature of the detector, injector, and oven were 200°C, 150°C and 100°C, respectively.

Total protein levels were determined on a plate reader (Bio-Tek Instruments, Winooski, VT) using a BCA-assay kit (Pierce, Rockford, IL) according to the manufacturer’s instructions. Total chlorophyll *a* was methanol extracted and quantified on an Olis DW2000 spectrophotometer (On-Line Instrument Systems, Inc., GA).

### *In vivo* ^15^N_2_ incorporation assay

All strains were grown under light/dark conditions as mentioned above, and 50 mL cultures were transferred into 125 mL airtight glass vials. 10 μM DCMU was added to the cultures, vials were flushed with pure N_2_, and cultures were incubated under light/dark conditions. Cells in the sealed vials were cultured overnight and at the time point D1, 8 mL of headspace gas was removed, followed by injection of 8 mL of ^15^N_2_ gas (98%^+^; Cambridge Isotope Laboratories, Inc.). After 8 h of incubation at 30°C in light (50 μmol photons · m^−2^ · s^−1^), the cultures were collected and dried in a laboratory oven at 50°C-60°C for 24 h. The dried pellets were ground, weighed, and sealed into tin capsules. Isotope ratios were measured by Elemental Analyzer-Isotope Ratio Mass Spectrometry (EA-IRMS, Thermo Fisher Scientific), and are showed as δ^15^N (‰), of which the number is a linear transform of the isotope ratios ^15^N/^14^N, representing the per mille difference between the isotope ratios in a sample and in atmospheric N_2_ (49). Data presented are mean values based on at least two biological replicate cultures.

### Western blot analysis

The coding region of the *nifD* gene (cce_0560) from Cyanothece 51142 was PCR-amplified from the genomic DNA of *Cyanothece* 51142, using the primers shown in Table S3. The PCR fragment was ligated into the expression vector pET28a cleaved by *Nde*I and *BamH*I. The resulting plasmid pET28a-nifD was used to produce the NifD protein with an N-terminal His_6_ tag. For overproduction of the NifD protein, *Escherichia coli* BL21 (DE3) was transformed with plasmid pET28a-nifD and cultivated in LB medium at 37 °C to an optical density at 600 nm (A_600_) of 0.3. Protein expression was induced by the addition of 0.2 mM isopropyl-*β-*D-thiogalactopyranoside, and the culture was incubated for another 18 h at 20 °C. After the cells were harvested, purification of NifD by nickel-nitrilotriacetic acid affinity chromatography was performed. Briefly, harvested *E. coli* cells were resuspended in 20 mM HEPES buffer (pH 7.0) containing 100 mM NaCl, and 2 mM *β*-mercaptoethanol supplemented with a protease inhibitor cocktail (Sigma-Aldrich). Lysozyme was added to a concentration of 1 mg/ml, and the cells were lysed by freezing-thawing, followed by sonication. After cells were centrifuged at 13,000 rpm, Tris-HCl buffer (pH 8.0) was added to the supernatant (final concentration, 50 mM), and it was loaded onto a Ni-NTA agarose column (0.2 ml). After bound proteins were washed with the starting buffer containing 1 M NaCl, they were eluted with 0.3 ml of the starting buffer containing 250 mM imidazole. The purified protein was stored at -20 °C and used as the positive control for western blot assay.

Cyanobacterial cells cultured in N-free medium under L/D cycles were collected at time point D4 (4 hours after dark phase) and resuspended in 0.5 mL TG buffer (10 mM Tris-HCl [pH 8.0], 10% glycerol) containing a protease inhibitor cocktail (Sigma-Aldrich). A 0.5 mL volume of sterilized, acid-washed glass beads was added to the cells, and the mixture was disrupted using a bead beater (BioSpec Products). The resultant mixture was centrifuged for 10 min at 7,500 × g, and the supernatant was transferred into a new tube. The amount of protein was determined using a bicinchoninic acid (BCA) protein assay reagent (Thermo Scientific).

Fifteen μg of total protein extract from each sample was solubilized with 8 × sample buffer (10 ml of 0.5 M Tris [pH 6.8], 15 ml of 70% glycerol, 8 ml of 20% sodium dodecyl sulfate, 4 ml of *β*-mercaptoethanol, 4 ml of 0.1% bromophenol blue) at 70 °C for 10 min and fractionated on a sodium dodecyl sulfate (0.1%, wt/vol)-polyacrylamide (12.5%, wt/vol) gel by electrophoresis. After electrophoresis, proteins were eletrotransferred to a polyvinylidene fluoride (PVDF) membrane (Millipore), blocked with 5% bovine serum albumin (BSA) for 2 hours at room temperature, and then incubated with the primary antibodies against the NifD protein from *Rhodospirillum rubrum* (50, 51) diluted in 1.5% BSA (1:2,000) overnight at 4 °C. The HRP-conjugated secondary antibody Goat Anti-Rabbit IgG (H + L)-HRP Conjugate (Bio-Rad) was diluted at 1:5,000 in 1.5% BSA. Immunodetection was performed using Western blotting Luminol Reagent (Millipore).

## Acknowledgements

This study was supported by the National Science Foundation (MCB-1331194). We thank Prof. Huimin Zhao and his research group (University of Illinois) for introducing us to the use of the DNA Assembler method; Lingxia Zhao, Xiujun Duan, and Alicia Lohman for expert technical assistance and members of the research groups of Himadri Pakrasi, Costas Maranas, Tae Seok Moon, and Fuzhong Zhang for critical scientific discussions. D.L., M.L., J.Y., H.B.P. and M.B.P. designed the experiments; D.L., M.L., J.Y. and M.B.P. performed the experiments; D.L., M.L., H.B.P. and M.B.P. wrote the paper.

## REFERENCES

1. Stokstad E. 2016. The nitrogen fix. Science 353:1225–1227.

2. Good A. 2018. Toward nitrogen-fixing plants. Science 359:869–870.

3. Vicente EJ, Dean DR. 2017. Keeping the nitrogen-fixation dream alive. Proc Natl Acad Sci U S A 114:3009–3011.

4. Buren S, Rubio LM. 2018. State of the art in eukaryotic nitrogenase engineering. FEMS Microbiol Lett 365:fnx274.

5. Curatti L, Rubio LM. 2014. Challenges to develop nitrogen-fixing cereals by direct *nif*-gene transfer. Plant Sci 225:130–137.

6. Dos Santos PC, Fang Z, Mason SW, Setubal JC, Dixon R. 2012. Distribution of nitrogen fixation and nitrogenase-like sequences amongst microbial genomes. BMC Genomics 13:162.

7. Hu Y, Ribbe MW. 2015. Nitrogenase and homologs. J Biol Inorg Chem 20:435- 445.

8. Mus F, Alleman AB, Pence N, Seefeldt LC, Peters JW. 2018. Exploring the alternatives of biological nitrogen fixation. Metallomics 10:523–538.

9. Hu Y, Ribbe MW. 2013. Nitrogenase assembly. Biochim Biophys Acta 1827:1112- 1122.

10. Sickerman NS, Ribbe MW, Hu Y. 2017. Nitrogenase cofactor assembly: an elemental inventory. Acc Chem Res 50:2834–2841.

11. Sickerman NS, Rettberg LA, Lee CC, Hu Y, Ribbe MW. 2017. Cluster assembly in nitrogenase. Essays Biochem 61:271–279.

12. Sickerman NS, Hu Y, Ribbe MW. 2017. Nitrogenase assembly: strategies and procedures. Methods Enzymol 595:261–302.

13. Dixon RA, Postgate JR. 1972. Genetic transfer of nitrogen fixation from *Klebsiella pneumoniae* to *Escherichia coli*. Nature 237:102–103.

14. Wang L, Zhang L, Liu Z, Zhao D, Liu X, Zhang B, Xie J, Hong Y, Li P, Chen S, Dixon R, Li J. 2013. A minimal nitrogen fixation gene cluster from *Paenibacillus* sp. WLY78 enables expression of active nitrogenase in *Escherichia coli*. PLoS Genet 9:e1003865.

15. Yang J, Xie X, Wang X, Dixon R, Wang YP. 2014. Reconstruction and minimal gene requirements for the alternative iron-only nitrogenase in *Escherichia coli*. Proc Natl Acad Sci U S A 111:E3718-3725.

16. Smanski MJ, Bhatia S, Zhao D, Park Y, L BAW, Giannoukos G, Ciulla D, Busby M, Calderon J, Nicol R, Gordon DB, Densmore D, Voigt CA. 2014. Functional optimization of gene clusters by combinatorial design and assembly. Nat Biotechnol 32:1241–1249.

17. Yang J, Xie X, Yang M, Dixon R, Wang YP. 2017. Modular electron-transport chains from eukaryotic organelles function to support nitrogenase activity. Proc Natl Acad Sci U S A 114:E2460-E2465.

18. Cheng Q, Day A, Dowson-Day M, Shen GF, Dixon R. 2005. The *Klebsiella pneumoniae* nitrogenase Fe protein gene (*nifH*) functionally substitutes for the *chlL* gene in *Chlamydomonas reinhardtii*. Biochem Biophys Res Commun 329:966–975.

19. Zamir A, Maina CV, Fink GR, Szalay AA. 1981. Stable chromosomal integration of the entire nitrogen fixation gene cluster from *Klebsiella pneumoniae* in yeast. Proc Natl Acad Sci U S A 78:3496–3500.

20. Lopez-Torrejon G, Jimenez-Vicente E, Buesa JM, Hernandez JA, Verma HK, Rubio LM. 2016. Expression of a functional oxygen-labile nitrogenase component in the mitochondrial matrix of aerobically grown yeast. Nat Commun 7:11426.

21. Perez-Gonzalez A, Kniewel R, Veldhuizen M, Verma HK, Navarro-Rodriguez M, Rubio LM, Caro E. 2017. Adaptation of the GoldenBraid modular cloning system and creation of a toolkit for the expression of heterologous proteins in yeast mitochondria. BMC Biotechnol 17:80.

22. Buren S, Young EM, Sweeny EA, Lopez-Torrejon G, Veldhuizen M, Voigt CA, Rubio LM. 2017. Formation of nitrogenase NifDK tetramers in the mitochondria of *Saccharomyces cerevisiae*. ACS Synth Biol 6:1043–1055.

23. Allen RS, Tilbrook K, Warden AC, Campbell PC, Rolland V, Singh SP, Wood CC. 2017. Expression of 16 nitrogenase proteins within the plant mitochondrial matrix. Front Plant Sci 8:287.

24. Ivleva NB, Groat J, Staub JM, Stephens M. 2016. Expression of active subunit of nitrogenase via integration into plant organelle genome. PLoS One 11:e0160951.

25. Oldroyd GE, Dixon R. 2014. Biotechnological solutions to the nitrogen problem. Curr Opin Biotechnol 26:19–24.

26. Falcon LI, Magallon S, Castillo A. 2010. Dating the cyanobacterial ancestor of the chloroplast. ISME J 4:777–783.

27. Stockel J, Welsh EA, Liberton M, Kunnvakkam R, Aurora R, Pakrasi HB. 2008. Global transcriptomic analysis of *Cyanothece* 51142 reveals robust diurnal oscillation of central metabolic processes. Proc Natl Acad Sci U S A 105:6156‱6161.

28. Cerveny J, Sinetova MA, Valledor L, Sherman LA, Nedbal L. 2013. Ultradian metabolic rhythm in the diazotrophic cyanobacterium *Cyanothece* sp. ATCC 51142. Proc Natl Acad Sci U S A 110:13210–13215.

29. Bandyopadhyay A, Elvitigala T, Welsh E, Stockel J, Liberton M, Min H, Sherman LA, Pakrasi HB. 2011. Novel metabolic attributes of the genus *Cyanothece*, comprising a group of unicellular nitrogen-fixing cyanobacteria. mBio 2:e00214.

30. Bandyopadhyay A, Elvitigala T, Liberton M, Pakrasi HB. 2013. Variations in the rhythms of respiration and nitrogen fixation in members of the unicellular diazotrophic cyanobacterial genus *Cyanothece*. Plant Physiol 161:1334–1346.

31. Shih PM, Wu D, Latifi A, Axen SD, Fewer DP, Talla E, Calteau A, Cai F, Tandeau de Marsac N, Rippka R, Herdman M, Sivonen K, Coursin T, Laurent T, Goodwin L, Nolan M, Davenport KW, Han CS, Rubin EM, Eisen JA, Woyke T, Gugger M, Kerfeld CA. 2013. Improving the coverage of the cyanobacterial phylum using diversity-driven genome sequencing. Proc Natl Acad Sci U S A 110:1053–1058.

32. Shao Z, Zhao H, Zhao H. 2009. DNA assembler, an *in vivo* genetic method for rapid construction of biochemical pathways. Nucleic Acids Res 37:e16.

33. Taton A, Unglaub F, Wright NE, Zeng WY, Paz-Yepes J, Brahamsha B, Palenik B, Peterson TC, Haerizadeh F, Golden SS, Golden JW. 2014. Broad-host-range vector system for synthetic biology and biotechnology in cyanobacteria. Nucleic Acids Res 42:e136.

34. Temme K, Zhao D, Voigt CA. 2012. Refactoring the nitrogen fixation gene cluster from *Klebsiella oxytoca*. Proc Natl Acad Sci U S A 109:7085–7090.

35. Gosink MM, Franklin NM, Roberts GP. 1990. The product of the *Klebsiella pneumoniae nifX* gene is a negative regulator of the nitrogen fixation (*nif*) regulon. J Bacteriol 172:1441–1447.

36. Ng AH, Berla BM, Pakrasi HB. 2015. Fine-tuning of photoautotrophic protein production by combining promoters and neutral sites in the cyanobacterium *Synechocystis* sp. Strain PCC 6803. Appl Environ Microbiol 81:6857–6863.

37. Liu D, Pakrasi HB. 2018. Exploring native genetic elements as plug-in tools for synthetic biology in the cyanobacterium *Synechocystis* sp. PCC 6803. Microb Cell Fact 17:48.

38. Tamagnini P, Leitao E, Oliveira P, Ferreira D, Pinto F, Harris DJ, Heidorn T, Lindblad P. 2007. Cyanobacterial hydrogenases: diversity, regulation and applications. FEMS Microbiol Rev 31:692–720.

39. Zhang X, Sherman DM, Sherman LA. 2014. The uptake hydrogenase in the unicellular diazotrophic cyanobacterium *Cyanothece* sp. strain PCC 7822 protects nitrogenase from oxygen toxicity. J Bacteriol 196:840–849.

40. Wunschiers R, Batur M, Lindblad P. 2003. Presence and expression of hydrogenase specific C-terminal endopeptidases in cyanobacteria. BMC Microbiol 3:8.

41. Rippka R, Deruelles J, Waterbury JB, Herdman M, Stanier RY. 1979. Generic assignments, strain histories and properties of pure cultures of cyanobacteria. J Gen Microbiol 111:1–61.

42. Bergkessel M, Guthrie C. 2013. Chemical transformation of yeast. Methods Enzymol 529:311–320.

43. Gibson DG, Young L, Chuang RY, Venter JC, Hutchison CA 3rd, Smith HO. 2009. Enzymatic assembly of DNA molecules up to several hundred kilobases. Nat Methods 6:343–345.

44. Golden SS, Brusslan J, Haselkorn R. 1987. Genetic engineering of the cyanobacterial chromosome. Methods Enzymol 153:215–231.

45. Williams JGK. 1988. Construction of specific mutations in photosystem II photosynthetic reaction center by genetic engineering methods in *Synechocystis* 6803, p 766–778, Methods Enzymol, vol Volume 167. Academic Press.

46. Kruse O, Rupprecht J, Mussgnug JH, Dismukes GC, Hankamer B. 2005. Photosynthesis: a blueprint for solar energy capture and biohydrogen production technologies. Photochem Photobiol Sci 4:957–970.

47. Oda Y, Samanta SK, Rey FE, Wu L, Liu X, Yan T, Zhou J, Harwood CS. 2005. Functional genomic analysis of three nitrogenase isozymes in the photosynthetic bacterium *Rhodopseudomonas palustris*. J Bacteriol 187:7784–7794.

48. Bandyopadhyay A, Stockel J, Min H, Sherman LA, Pakrasi HB. 2010. High rates of photobiological H_2_ production by a cyanobacterium under aerobic conditions. Nat Commun 1:139.

49. Montoya JP, Voss M, Kahler P, Capone DG. 1996. A simple, high-precision, high-sensitivity tracer assay for N_2_ fixation. Appl Environ Microbiol 62:986–993.

50. Colon-Lopez MS, Sherman DM, Sherman LA. 1997. Transcriptional and translational regulation of nitrogenase in light-dark- and continuous-light-grown cultures of the unicellular cyanobacterium *Cyanothece* sp. strain ATCC 51142. J Bacteriol 179:4319–4327.

51. Grunwald SK, Lies DP, Roberts GP, Ludden PW. 1995. Posttranslational regulation of nitrogenase in *Rhodospirillum rubrum* strains overexpressing the regulatory enzymes dinitrogenase reductase ADP-ribosyltransferase and dinitrogenase reductase activating glycohydrolase. J Bacteriol 177:628–635.

52. Curatti L, Brown CS, Ludden PW, Rubio LM. 2005. Genes required for rapid expression of nitrogenase activity in *Azotobacter vinelandii*. Proc Natl Acad Sci U S A 102:6291–6296.

53. Han Y, Lu N, Chen Q, Zhan Y, Liu W, Lu W, Zhu B, Lin M, Yang Z, Yan Y. 2015. Interspecies transfer and regulation of *Pseudomonas* stutzeri A1501 nitrogenfixation island in *Escherichia coli*. J Microbiol Biotechnol 25:1339–1348.

54. Kavanagh EP, Hill S. 1993. Oxygen inhibition of nitrogenase activity in *Klebsiella pneumoniae*. J Gen Microbiol 139:1307–1314.

